# A scalable architecture for tuning multistate differentiation ratios in synthetic microbial consortia

**DOI:** 10.64898/2025.12.17.694810

**Authors:** Chloé Sasson, Julien Capin, Quentin Boussau, Amanda Abi-Khalil, Mélanie Guyot, Capucine Mayoud, Anna-Sophie Fiston-Lavier, Elsa Fristot, Judith Bruyère, Xavier Devos, Martin Cohen-Gonsaud, Diego Cattoni, Jerome Bonnet

## Abstract

Establishing synthetic microbial consortia in competitive environments is often compromised by stochastic colonization bottlenecks, where founder effects lead to the unpredictable dominance of a single strain. Here, we overcome this challenge by engineering a differentiation abacus, a scalable, single-layer recombinase architecture that enables a single progenitor cell to differentiate into up to twelve distinct subpopulations. By arranging competitive excision sites in a linear array, we demonstrate that differentiation ratios can be programmed through rationally tuning recombination-site kinetics and inter-site spacing. This architecture allows the generation of strictly mutually exclusive phenotypes with tunable composition, scaling from simple two-state systems to complex multi-state ensembles without the need for multilayered regulation. Finally, we validate the system’s utility in a mouse tumor model, showing that *in situ* differentiation establishes robust, homogeneous consortia that overcome the colonization variability associated with pre-assembled mixtures. This work provides a versatile and scalable framework for reliably controlling consortia composition for bioproduction, synthetic ecology, and engineered living therapies.

## INTRODUCTION

An overall goal of synthetic biology is to reprogram cellular behavior for applications in biomanufacturing, medicine, and the environment^1–3^. Over the past decades, researchers have reprogrammed living cells to encode Boolean logic^4,5^, memory^6–8^, robust oscillations^9,10^, and spatial pattern formation^11,12^. As cellular programs grow in complexity, dividing labor across a consortium of multiple cell types offers a powerful strategy to implement sophisticated behavior at the population level while reducing metabolic burden on individual cells and improving the overall system’s robustness^13,14^.

Engineered microbial consortia can be obtained by preassembling multiple strains, often using stabilization strategies such as synthetic metabolic dependencies and intercellular communication systems^15,16^. However, preassembled consortia face significant limitations, such as difficulty accessing different strain ratios once in the environment and limited practical scalability to a large number of phenotypes. Perhaps most critically, pre-assembled consortia might struggle to achieve reliable co-colonization of specific confined niches. For example, cancer therapies benefit from combination approaches^17^. Bacterial cancer therapy offers a unique opportunity to achieve this locally, as engineered strains can colonize tumors and deliver multiple biotherapeutics *in situ*. However, engineering a single strain to express multiple effectors imposes a substantial metabolic burden on the cell, which typically reduces secretion efficiency and therapeutic output^18^. Furthermore, delivering pre-assembled consortia of specialized strains, each producing a different effector, has shown limited efficacy due to poor tumor colonization when administered intravenously^19^. This bottleneck can generate a “founder” or “priority” effect^20–22^, in which the first strain to successfully establish often expands to dominance and competitively excludes the others, making it challenging to ensure that all members of the consortium are represented at the tumor site and reproduce the intended proportions for efficient therapeutic outcome^23^.

To overcome these bottlenecks, biological systems have evolved to form consortia with distinct functions and precise ratios through cellular differentiation. From the development of multicellular tissues^24^ to division of labor and bet-hedging in bacterial populations^25,26^, complex multicellular systems often arise from a single progenitor lineage that diversifies *in situ*. Emulating this strategy would allow a single engineered strain to colonize a target niche at low burden before generating *in situ* a therapeutic consortium. Realizing this approach requires synthetic tools that provide tunable multi-state control. Researchers have begun engineering single-cell bacterial differentiation systems using plasmid segregation mechanisms that rely on asymmetric inheritance of engineered plasmids during cell division to generate two subpopulations with distinct phenotypes^27,28^. However, these designs are typically restricted to binary cell-fate decisions and offer limited control over population ratios.

Serine integrases enable synthetic differentiation via irreversible DNA excision or inversion and have been widely used for genome engineering and for recombinase-based logic and memory in living cells^29^. These enzymes were used to implement robust bistable fate decisions, including architectures in which differentiation is coupled to survival^30^, and related strategies such as synthetic phase variation can stochastically switch between two phenotypes, while higher-order states require additional recombinases^31^. However, these approaches are limited to two-state differentiation and do not readily scale to many differentiated phenotypes with tunable population ratios. More complex recombinase frameworks can expand diversity (*e.g.*, Brainbow^32^) or encode history-dependent programs^33,34^, but they are intrinsically heterogeneous or designed for multi-input logic rather than for control over phenotypic ratios. Recent multilayer branching circuits with tunable proportions^35^, primarily demonstrated in yeast, remain constrained in scalability by limited site orthogonality, more complex regulatory layers, and often yield mixed-output cells rather than mutually exclusive fates. This lack of strict mutual exclusivity undermines the metabolic division of labor and can reintroduce the physiological burden that synthetic consortia aim to eliminate. For applications that demand strict metabolic division of labor, it is therefore desirable to develop differentiation architectures that convert lineage diversity into clearly partitioned, mutually exclusive functional states at the single-cell level.

To our knowledge, no current strategy provides a generic, highly scalable differentiation system in bacteria that combines (i) a genetically simple, compact, single-layer architecture with a low-burden parental state, (ii) differentiation triggered by a single transient input, (iii) high scalability to many alternative phenotypes, (iv) strictly mutually exclusive output in each differentiated cell types, and (v) straightforward, designable tuning of the resulting phenotypic ratios. Such a system would allow a single strain to colonize or expand as an undifferentiated population and, upon induction, diversify into a controlled ensemble of specialized cell types that function together as a synthetic consortium, each assigned a specific single genetic expression task.

In this work, we engineer a scalable, single-layer recombinase differentiation system that enables a single bacterial progenitor cell to generate up to twelve distinct subpopulations in response to a single input. Our system is based on competitive excision of recombination sites flanking genes of interest and arranged in a linear competitive excision array, a configuration reminiscent of an abacus. Recombination results in gene shuffling and leads to mutually exclusive expression of different genes in each daughter cell. We first implement two- and three-state differentiation systems in which each state drives the expression of distinct fluorescent proteins and show that the relative proportions between daughter cells can be tuned by combining *att* sites with varying recombination efficiencies. We then demonstrate the scalability of our “differentiation abacus” by engineering six- and twelve-state differentiation systems by simply adding recombination sites along the DNA strand, without needing to redesign the basic circuit. Our architecture preserves a compact and uniform regulatory layer while still exploiting differences in *att*-site recombination efficiencies, allowing the number of differentiated bacterial phenotypes to scale in a predictable, designable manner. Finally, we establish a proof-of-concept in a colorectal tumor model in mice, demonstrating that, following colonization by progenitor cells, transient induction of differentiation with exogenous molecules generates a broader and more balanced spectrum of phenotypes than when preassembled consortia are injected. This work provides a foundation for reliably controlling the proportions and numbers of differentiated phenotypes in bacterial populations, with applications in bioproduction, bioremediation, morphogenetic engineering, and bacterial therapy.

## RESULTS

### Two- and Three-States Differentiation Circuit With Single Recombinase Input

We started by building a bacterial differentiation system based on irreversible, competitive recombination reactions between multiple attachment (*att*) sites catalyzed by serine integrases^36^. Serine integrases catalyze unidirectional, site-specific recombination between *attP* (for Phage) and *attB* (for Bacterial) attachment sites (**Supplementary Fig. 1**). After recombination, the original *attP* and *attB* sites are converted into *attL (Left)* and *attR (Right)*, respectively, preventing further recombination unless a Recombination Directionality Factor (RDF) is coexpressed with the integrase^8,37^. Without RDF, the reaction is irreversible, and the system acts as a *de facto* memory switch.

We first built a two-state differentiation bacterial system controlled by the Bxb1 integrase^38^ by designing a recombinase target construct in which a single *attP* site, preceded by a constitutive promoter, is followed by two identical *attB* sites, *attB1* and *attB2,* positioned in parallel orientations, resulting in excision when recombination occurs (**Fig. 1a**). We selected this specific arrangement of *att* sites based on an optimized design we previously characterized^39^. We placed a strong terminator upstream of *attB1*, a sequence containing a first gene of interest (GOI1) preceded by a ribosome binding site (RBS) between *attB1* and *attB*2, and a sequence containing a second gene of interest (GOI2). In state 0 (S0), the terminator blocks the RNA polymerase and the expression of downstream genes. Upon Bxb1 expression, competitive recombination reactions (*attP*/a*ttB1* or *attP/attB2)* result in the removal of the terminator, triggering GOI1 expression (state 1, S1) or in the removal of the terminator-*attB1*-GOI1 sequence, resulting in GOI2 expression (state 2, S2).

**Figure 1:**
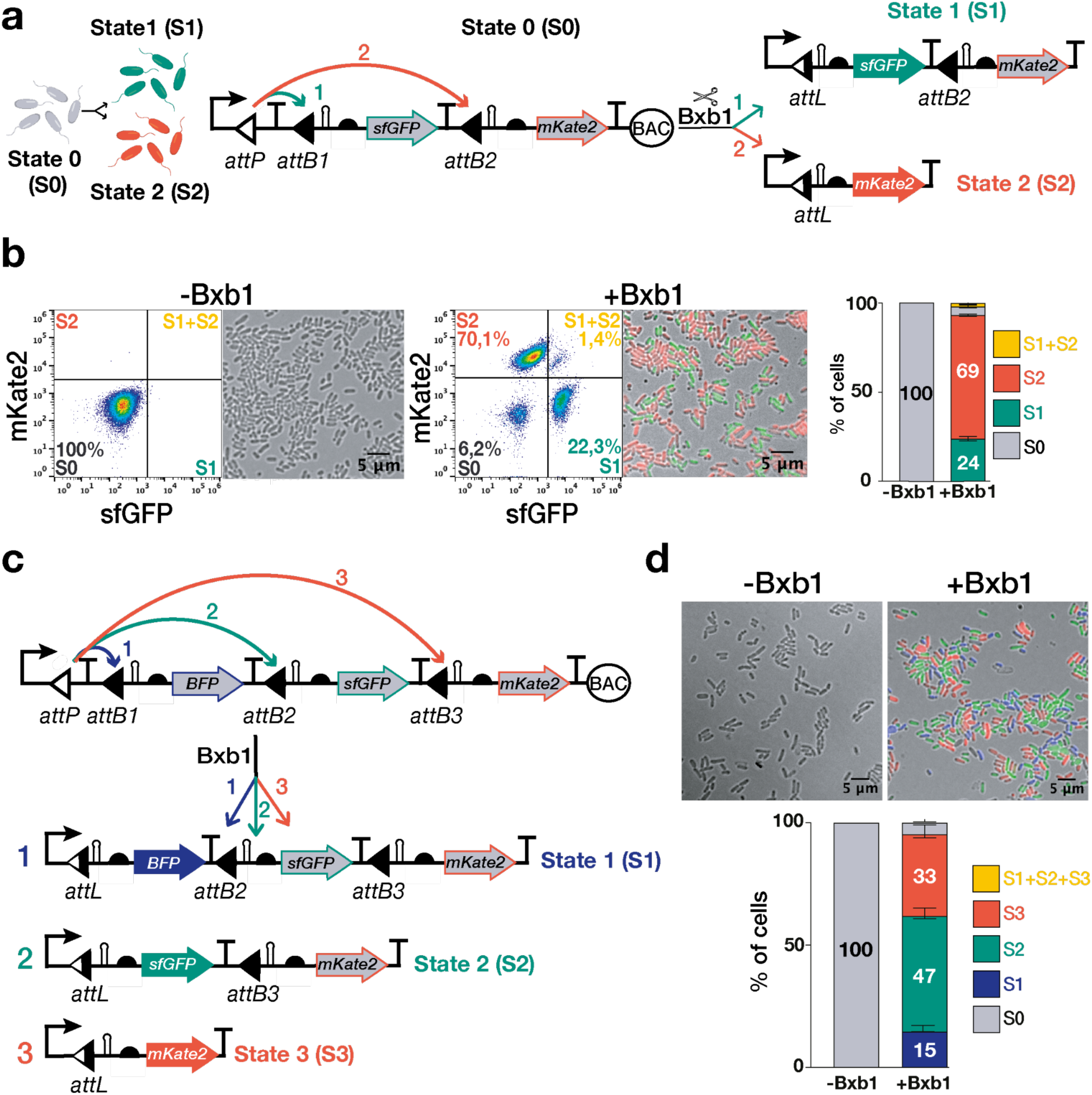
Two and three-state bacterial differentiation controlled by the Bxb1 recombinase. **a.** Architecture of the two-state differentiation system. In state 0 (S0), bacteria are undifferentiated and do not express any gene of interest. Upon Bxb1 expression, mutually exclusive excision reactions result in cells harboring one of the two possible recombination outcomes and expressing different genes of interest. Here, the outcomes are visualised by expression of the fluorescent proteins superfolder GFP (state 1, S1) and mKate2 (state 2, S2). **b.** Integrase-mediated two-state differentiation in living cells. Representative example of flow cytometry profiles and associated fluorescence microscopy images of bacteria harboring the two-state differentiation circuit in the absence (left panel) or presence (middle panel) of the Bxb1 recombinase. Right panel: Corresponding bar charts displaying the percentage of cells in each state. Bacteria were co-transformed with the target construct and a constitutive Bxb1 recombinase, diluted 1:100 into LB medium supplemented with the appropriate antibiotics (chloramphenicol and ampicillin), and grown for 16h at 30°C with shaking at 250 rpm. n=3 independent experiments performed on different days in triplicate, mean±SD. **c.** Architecture and recombination states of the three-state differentiation system. **d.** Integrase-mediated three-state differentiation in living cells, as in c.

To perform a quantitative analysis of recombination efficiency and dynamics of this two-state differentiation design, we built a recombination target that expresses either the superfolder Green Fluorescent Protein (sfGFP) in S1 or the red fluorescent protein mKate2 in S2. We monitored bacterial fate by flow cytometry and wide-field fluorescent microscopy. Each gene was preceded by a strong RBS and a self-cleaving ribozyme^40^, to remove any undesirable 5’ UTR resulting from different recombination reactions that could alter protein expression (**Fig. 1a**).

We co-transformed *E. coli NEB 10 Beta* cells with a plasmid bearing the two-state recombination target and a medium-copy plasmid constitutively expressing Bxb1. When employing a low-copy plasmid (pSC101, 5-10 copies/cell) for the target sequence, a fraction of the cells (∼16%) exhibited both states simultaneously (**Supplementary Fig. 2**). We hypothesized that having multiple copies of the target within a single progenitor cell could be causing daughter cells to harbour multiple states simultaneously. To confirm this hypothesis, we moved away from multiple-copy plasmids by either integrating a single copy of the recombination target into the bacterial chromosome or cloning the target into a bacterial artificial chromosome^41^ (BAC, ∼1 copy/cell). Chromosomal integration and BAC cloning produced single-state daughter cells as observed by flow cytometry. We decided to use the BAC backbone as it provides an easy-to-manipulate vector with equivalent chromosomal copy-number conditions^42^.

Detailed quantification of the population across each differentiated state revealed that, on average, 69% of cells were in state 2 and 24% in state 1, corresponding to a population ratio of ∼2.9:1 (*attB2*:*attB1*), indicating a preferential differentiation toward S2, and suggesting a difficulty for Bxb1 to excise sites at short DNA distances (**Fig. 1c**). Differentiation and proportions of states were further confirmed using time-lapse microscopy (**Supplementary Fig. 3-5**).

Finally, to demonstrate the versatility of the differentiation system across chassis, we chromosomally integrated the two-state differentiation circuit into the bacterial strain *E. coli Nissle 1917* (*EcN*), a chassis of therapeutic interest. The circuit was inserted at distinct genomic loci (*lacZ* for *EcN* vs *yfc* in *E. coli NEB 10 Beta).* Upon constitutive expression of the Bxb1 recombinase, the resulting state distributions between the two chassis were comparable, demonstrating that our differentiation system is robust across two bacterial hosts and two distinct genomic integration sites (**Supplementary Fig. 6**).

To demonstrate the versatility of our strategy, we also explored several other architectures, such as designs based on competitive inversion, (**Supplementary Fig. 7**), competing inversion and excision reactions (**Supplementary Fig. 8**), or designs using palindromic central dinucleotides^43^ in which the system loses directional asymmetry, enabling Bxb1 to catalyze both inversion and excision on the same pair of *att* sites (**Supplementary Fig. 9**). Inversion-based designs are of particular interest, as they allow the DNA sequence, if desired, to be reverted to its original state through the addition of an excisionase. All these designs successfully enabled functional cellular differentiation into two distinct phenotypes, though with varying phenotypic ratios. These differences likely reflect underlying biases toward specific reaction types or dinucleotide sequences.

We then sought to expand the system beyond two differentiated states. To this end, the abacus-like design provided a straightforward strategy for scaling higher-order states by simply concatenating additional *att* sites along the DNA strand. We thus introduced a third *attB* site and incorporated an additional reporter gene encoding the Blue Fluorescent Protein (BFP) (**Fig. 1d**). Upon expression of Bxb1, cells bearing the three-state differentiation systems differentiated into three phenotypically distinct populations in S1, S2, and S3, respectively expressing BFP, sfGFP, or mKate2 (**Fig. 1e)**. Consistent with our previous observations, we observed preferential excision at the distal sites (47% at the second, 33% at the third) compared to the proximal first site (15%). These results reinforced the idea that Bxb1 encounters a barrier to recombine the proximal site, likely due to steric constraints imposed by the short inter-site distance.

To distinguish between physical constraints and biological burden as the causes of this bias in differentiation ratios, we first assessed reporter-specific effects. We observed that mKate2-expressing strains exhibited lower fitness than sfGFP-expressing ones (**Supplementary Fig. 10**). After exchanging fluorescent reporters, the bias against the proximal *attB1* site persisted (**Supplementary Fig. 11**). Thus, while this differential metabolic burden might explain minor fluctuations in observed differentiation ratios, it could not account for the strong, persistent bias against the proximal *attB1* site. We therefore hypothesized that the short genomic distance between *attP* and *attB1* imposed a physical barrier to recombination.

By systematically increasing the spacer length from 149 bp to over 17 kb (**Supplementary Fig. 12a**), we observed a striking reversal in recombination preference. While distances below ∼200 bp strongly reduced recombination at *attB1*, extending the spacer beyond ∼400 bp shifted the bias toward the proximal site, eventually plateauing at ∼70% efficiency (**Supplementary Fig. 12b**). We confirmed this effect in the three-state architecture, finding that the insertion of the *luxCDABE* operon was sufficient to rescue recombination at the proximal site, raising the proportion of cells in S1 from 15% to 57% (**Supplementary Fig. 13**). This behavior implies that inter-site distances below ∼200 bp impose a mechanical barrier, likely attributable to the persistence length of double-stranded DNA (∼150 bp)^44,45^. At short scales, the energy required to bend DNA likely exceeds the recombinase binding energy^46,47^. Beyond this stiffness threshold (>400 bp), the system transitions to a diffusive search regime, in which the higher effective concentration of the proximal site dictates the reaction rate (see **Supplementary Note 1**).

Taken together, these results demonstrate that multi-state differentiation systems can be implemented simply and efficiently using recombinase-mediated competitive excision reactions. Furthermore, the abacus design allows straightforward scalability to higher-order states by concatenating additional recombination sites. To achieve more precise, programmable control over differentiation ratios, we next exploited attachment site variants with distinct recombination efficiencies.

### Precise and Tunable Control of Phenotypic Ratios Using Attachment Site Variants With Differential Recombination Efficiencies

Zhang *et al*. recently explored modulation of Bxb1 recombination kinetics through targeted modifications of the DNA attachment sites^48^. By engineering *attP* with combinations of nucleotide substitutions, they generated multiple variants exhibiting distinct recombination rates, later reapplied by An *et al.*^35^. We used two of these variants: an *attP_fast* version carrying the substitutions G30A, G33T, G37C, C39T, and G42A, and an *attP_slow* version containing the single-point mutation C39A (**Fig. 2a**). Since our initial two-state differentiation design included one *attP* site and two *attB* sites, we redesigned the system to feature one *attB* site and two *attP* sites. Using this new architecture, we tested seven combinations involving the wild-type *attP* (*attP_WT*), *attP_fast*, and *attP_slow* to evaluate their impact on recombination dynamics.

**Figure 2:**
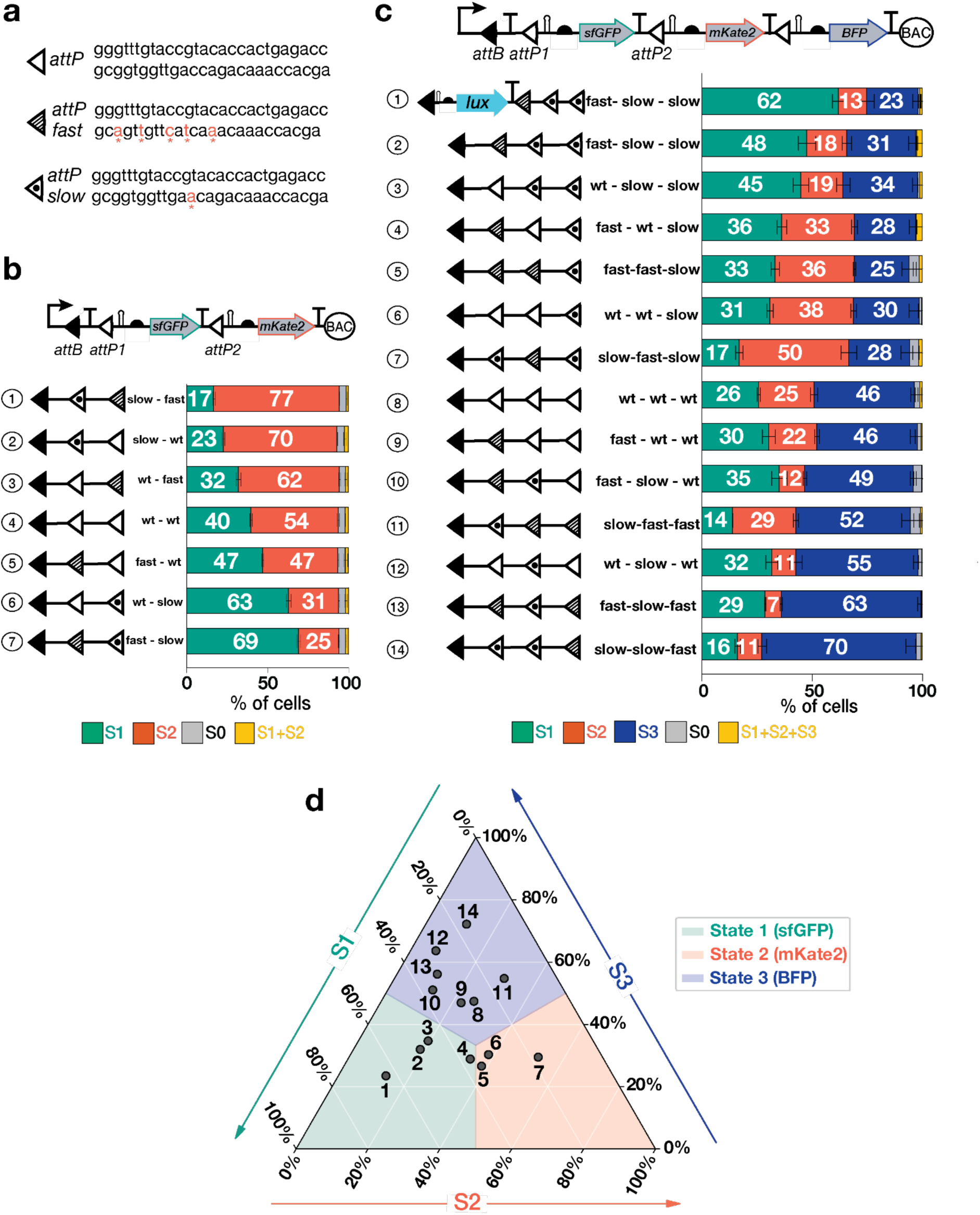
Tuning the proportion of differentiated cells using attachment site variants. **a.** DNA sequences of Bxb1 *attP* wild-type and two mutants, *attP_fast* and *attP_slow*. Mutated bases are shown in red with a star. **b.** Two-state differentiation circuits with different combinations of *attP* mutants (*attP_fast*, *attP_slow*, and *attP* corresponding to the wild-type) on a bacterial artificial chromosome (BAC), and their associated histograms obtained from flow cytometry measurements. **c.** Three-state differentiation circuits with different combinations of *attP* mutants (*attP_fast*, *attP_slow*, and *attP* corresponding to the wild-type) on a bacterial artificial chromosome (BAC), and their associated histograms obtained by flow cytometry. For two-state circuits, bacteria were grown for 16 hours at 30°C, for three-state circuits, bacteria were grown for 16 hours at 37°C, as BFP has a slower maturation rate^50^. **d.** Ternary plot visualizing the differentiation landscape accessible by the engineered three-state architectures. The axes represent the population fractions of states 1 (sfGFP), 2 (mKate2), and 3 (BFP). Numbered data points correspond to the specific circuit configurations listed in c.

Interestingly, modifying the circuit architecture to position two *attP* sites in competition for a single *attB* site significantly shifted differentiation ratios compared to the original design (two *attB* sites competing for one *attP*). While the genomic distances between sites remained constant, the population distribution shifted from a distinct 2.9:1 preference (state two dominant) to a more balanced 1.3:1 ratio (**Fig. 2b**). As the inter-site spacing (∼200 bp) is below the stiffness threshold, loop formation becomes critically sensitive to helical phasing. Consequently, the nine bp difference between the central dinucleotide cutting sites of the two architectures likely alters the rotational alignment of the asymmetric recombinase complex^49^. In this constrained topological context, we hypothesize that such phase shifts would differentially modulate the efficiency of synaptic complex assembly, thereby driving the observed change in recombination bias (**Supplementary Note 1**).

We next constructed a library of two-state differentiation circuits using attachment-site variants with distinct recombination kinetics. Consistent with their differential reaction rates, incorporating the *attP_fast* mutant increased the proportion of the corresponding differentiated state, while the *attP_slow* mutant reduced it. This combinatorial approach yielded a tunable range of population ratios. Specifically, the fast-slow configuration maximized the S1 population (69%) against S2 (25%), whereas the inverse slow-fast arrangement minimized S1 (17%) in favor of a dominant S2 population (77%), with the wild-type pair (wt-wt) yielding an intermediate distribution of 40% S1 and 54% S2 cells. (**Fig. 2b**).

We validated the scalability of this strategy by applying it to a three-state circuit, successfully generating a multiphenotypic consortium with precise ratio control (**Fig. 2c**). By varying site combinations, we shifted the population distribution to favor specific outcomes: a majority of cells in S1 (45%) when using a fast-slow-slow architecture; a majority of cells in S2 (52%) when using a slow-fast-slow architecture; and a majority of cells in S3 (70%) for the slow-slow-fast architecture.. Interestingly, switching from a wt-slow-slow to a fast-slow-slow configuration did not drastically increase the S1 population ratio (42 to 45%). Reaching a larger majority of S1 cells (62%) required a synergistic approach that combined a ‘fast’ variant (fast-slow-slow configuration) with the *lux* operon spacer to simultaneously increase recombinase affinity and relieve the DNA stiffness inhibiting recombination at the first site. Because the system naturally biases toward S3 (45% in the wild-type baseline), we found that the design rule of placing a ‘slow’ variant at the last position, at the condition that the efficiency of the two other proximal sites was not diminished, consistently counteracted this bias, redistributing the population into near-equal proportions (constructs 4, 5, and 6). We mapped the population compositions of all three-state architecture variants onto a ternary plot (**Fig. 2d**), revealing a comprehensive coverage of the differentiation space. By rationally permuting site kinetics and spacing, we could bias the consortium composition toward dominance by any of the three phenotypes. This broad phenotypic accessibility confirms the tunability of our “differentiation abacus”, showing its capability to generate diverse, reproducible consortia configurations from a single genetic architecture.

These findings establish a generalized framework for programming cellular differentiation ratios, enabling the rational construction of multi-state consortia with user-defined population architectures.

### High-Order Multistate Differentiation

We next aimed at testing the scalability of our design. To further expand the number of differentiation states, we started from the *attP*-*attB1*-*attB2* architecture and introduced additional *attB* sites, designing a system capable of supporting up to six distinct excision states (S1-S6). Due to the challenges of simultaneously detecting and quantifying seven distinct populations using a gene reporter or gene expression approach, we turned to long-read sequencing to estimate the proportions of the different recombination products. To this end, random stuffer DNA sequences of around 200 bp were inserted between the *attB* sites (**Fig. 3a**). A plasmid containing the differentiation array was then transformed into an *E. coli* strain constitutively expressing Bxb1 from the chromosome. Following transformation and overnight culture, plasmids containing recombined states were extracted and sequenced via long-read sequencing (**Fig. 3b**).

**Figure 3:**
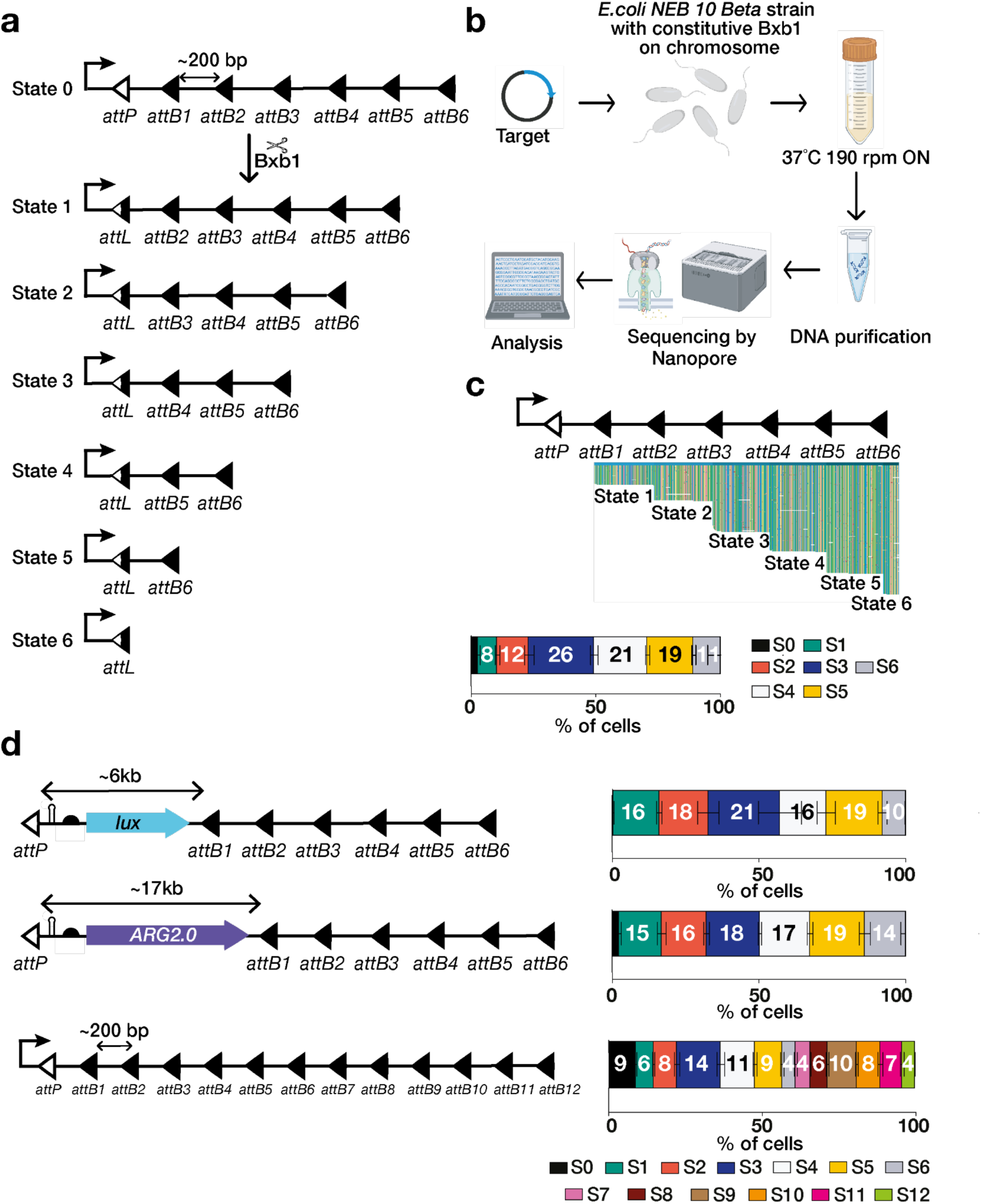
Recombinase-based one-input bacterial six- and twelve-state differentiation system by competitive excision. **a.** Schematic representation of the six-state circuit with the six recombined plasmids after Bxb1 excision. **b.** *E. coli NEB 10 Beta cells* carrying a chromosomally integrated constitutive Bxb1 cassette were transformed with the target plasmid and cultured overnight at 37 °C with shaking at 190 rpm. The following morning, DNA was extracted from the culture and subjected to long-read sequencing. The resulting reads were aligned to the target sequence and analyzed. **c.** Long-read sequencing analysis of the six-state differentiation system. Top: Visualization of single-molecule Nanopore reads aligned to the target locus, processed using the custom Python script RecombCompet (see methods). Distinct read patterns correspond to specific excision states. Bottom: Bar graph quantifying the relative abundance of each differentiated state in the population. Panel created with BioRender.com. **d.** Long-read sequencing on constructs modified with large genetic spacers (*lux*, ∼6 kb; *ARG2.0*, ∼17 kb) or expanded to twelve recombination sites, with their corresponding bar charts and population distribution profiles. For all experiments, n=3 independent experiments, mean ± SD.

Sequencing confirmed that Bxb1-mediated recombination successfully generated six distinct differentiated DNA states, demonstrating the system’s scalability (**Fig. 3c**). We quantified differentiation ratios and found that the first excision state (State 1), which requires recombination between *attP* and the most proximal *attB* site (∼200 bp distance), was the least frequent, representing only 8% of the population. This is consistent with our previous observation that short DNA loops are energetically unfavorable for recombinase synapsis (**Supplementary Fig. 12**). Conversely, State 3 was the most abundant (26%), followed by States 4 (21%) and 5 (19%). The final excision product, State 6, which requires recombination with the most distal site, comprised 11% of the population. We next sought to achieve more balanced state ratios. Based on our earlier findings that distance affects recombination efficiency, we inserted large DNA sequences, the *luxCDABE* operon or the acoustic reporter operon *ARG2.0*^51^, both generally used for visualizing bacteria *in vivo*, into the first position of the six-state differentiation construct (**Fig. 4d**). In these designs, we observed that all six differentiated states were represented at approximately equal proportions, around 16%. Read counts for each state are available in **Supplementary Fig. 14**. While multiple repeats could lead to sequence instability via recombination or replication slippage, we did not observe such an effect on the six-state recombination target after growing cells for ∼90 generations, even in *E. coli Nissle 1917*, which is RecA-positive (**Supplementary Fig. 15**). The stability is probably due to the low copy number of the construct.

**Figure 4:**
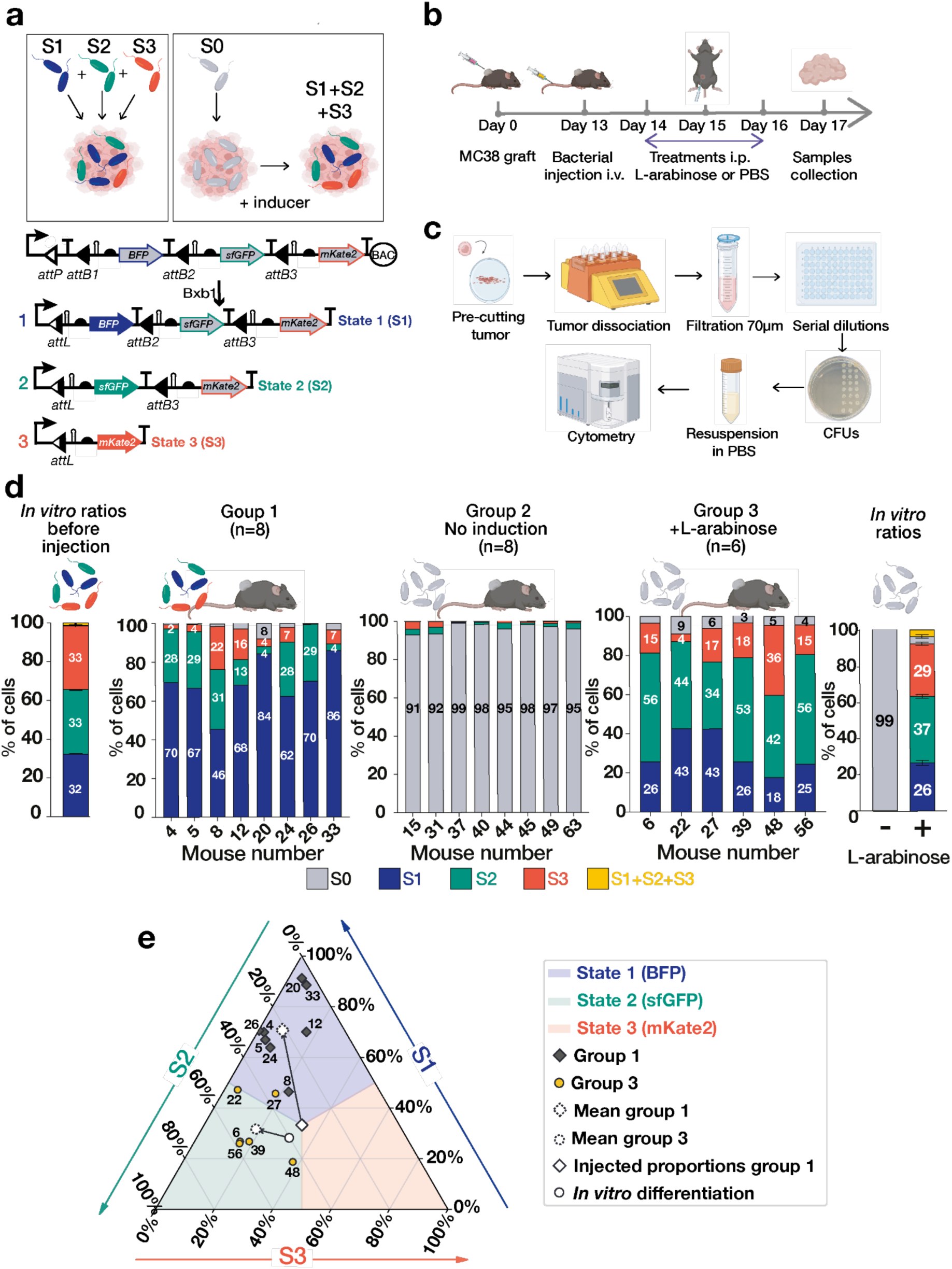
Three-state differentiation in a mouse tumor model. **a.** <u>Upper panel:</u> Experimental schematic comparing a three-state differentiation circuit with a control condition consisting of a mixture of three single-state strains in a murine cancer model. <u>Lower panel:</u> Genetic architecture and recombination states of the three-state differentiation circuit integrated into the *E. coli Nissle* chromosome. **b.** *In vivo* experimental workflow. MC38 colorectal cancer cells were implanted subcutaneously into the flanks of C57BL/6 mice. When tumours reached 150–200 mm³ (approximately 13 days post-implantation), 5 × 10⁶ bacteria were administered intravenously. Beginning one day after bacterial injection, mice received intraperitoneal injections of L-arabinose (1 g kg⁻¹) or PBS for three consecutive days. Tumours were harvested five days after bacterial colonization. **c.** Post-harvest tumour processing, including tissue dissociation, colony-forming unit (CFU) plating, and flow cytometry analysis. **d.** Flow cytometry–derived bar plots showing fluorescence-state distributions for individual mice in each experimental group. Numbers indicate individual mice. n = 8 for each group. Mice that were not colonized, as assessed by CFU plating (two in group 3), were excluded from the analysis. **e.** Ternary plot showing the relative proportions of S1, S2, and S3 for each mouse. Each point represents an individual mouse. Grey diamonds indicate group 1 (mixture of three single-state strains), and yellow circles indicate group 3 (three-state differentiation circuit with induction). The white diamond indicates the expected distribution for group 1 based on an initial 1:1:1 mixture (33% S1, 33% S2, 33% S3); the dashed white diamond indicates the group mean. The white circle indicates the expected distribution for group 3 (*in vitro* values), with the dashed white circle indicating the group mean. Arrows indicate deviation between expected and observed *in vivo* distributions. Panels a, b, c, and d were partially created with BioRender.com.

Finally, to demonstrate the high-order scalability of our architecture, we expanded the array to include twelve *attB* sites separated by ∼200 bp spacers (**Fig. 4d, bottom panel**). Long-read sequencing analysis confirmed that recombination successfully generated a highly diversified population containing every state. The differentiation profile showed individual subpopulations ranging from 4% to 14%, with a relatively significant fraction of undifferentiated cells (9%, S0). These data demonstrate that our competitive excision architecture can scale to generate at least 12 distinct phenotypes by concatenating additional att sites in response to a single input.

### Establishing Robust Multi-Phenotypic Consortia Inside Tumors

To evaluate the robustness of our differentiation system in complex physiological environments, we applied it to bacterial cancer therapy, an approach in which engineered bacteria must colonize tumors and deliver therapeutic payloads^19,52,53^. While combination therapies are highly desirable, expressing multiple payloads within a single strain imposes a metabolic burden that rapidly drives counter-selection, as exemplified by the fitness loss and slowdown in growth rate of bacteria expressing three fluorescent proteins compared to those expressing only one (**Supplementary Fig. 16**). Consequently, the field has largely relied on co-administering mixtures of specialized strains. While this approach can be successful via direct intratumoral administration^54,55^, it becomes more challenging when the intravenous (IV) route is used, as circulating bacteria must survive immune clearance, overcome vascular barriers to colonize the tumor, and compete for scarce ecological niches in an inflamed environment^19^. Recent work in a mouse tumor model showed that only a few clones reach the tumor and compete for growth, resulting in highly heterogeneous distribution within and across tumors^23^. Consequently, establishing robust IV-mediated co-colonization of tumors by multiple strains, particularly at defined ratios, remains a significant challenge in the field.

To quantify these colonization barriers in our model, we performed a pilot experiment using the probiotic *E. coli Nissle 1917* (*EcN*), a commonly used chassis for bacterial cancer therapies. We intravenously administered a 1:1 mixture of two *EcN* strains (each expressing a distinct fluorescent protein) to a small cohort of immunocompetent C57BL6/J mice bearing subcutaneous MC38 colorectal tumors. Analysis of colony-forming units (CFUs) and flow cytometry of bacteria recovered from dissociated tumors revealed that while single strains colonized tumors efficiently, co-colonization by mixed strains was highly heterogeneous across individuals (**Supplementary Fig. 17**). Furthermore, when performing in another cohort sequential injections of the two strains separated by several days, we observed that tumors were consistently colonized by either the first or the second strain, but rarely by both. These findings suggest a competitive-exclusion mechanism: successful colonization by the second strain occurs only when the first fails to establish, whereas poor colonization by the second correlates with robust pre-existing colonization by the first. We hypothesize that the tumor niche, once occupied, resists displacement by a subsequently introduced strain. Taken together, these results suggest, as already proposed by others, that multi-strain tumor colonization, even in this favorable model of transplanted tumors, is poorly reproducible and generally inefficient, thereby limiting the feasibility of this approach.

We reasoned that using our programmable differentiation system could address this issue: undifferentiated bacteria that initially colonize the tumor would provide a phenotypically homogeneous population, which could then be induced to differentiate into distinct subpopulations, each producing a different effector with controllable proportions. We thus sought to compare the two strategies for establishing a multi-strain consortium in tumors: co-injection of a premixed strain suspension comprising the various consortium members versus injection of a single progenitor strain that would differentiate and form the consortium *in situ*.

We integrated the three-state differentiation system at the *lacZ* locus on *EcN*. By performing *in vitro* recombination reactions, we isolated recombined daughter cells for each state, which we used to produce a preassembled consortia. We tested three experimental conditions in the same mouse colorectal tumor model (**Fig. 4a**). The first group of mice received a pre-assembled consortium containing an equal mixture of three *EcN* strains in differentiated states 1, 2, and 3. The remaining two groups were inoculated with *EcN* carrying the chromosomal three-state differentiation circuit and a plasmid bearing the arabinose-inducible Bxb1 recombinase. Starting one day post-injection, one group was treated with 1 g/kg L-arabinose intraperitoneally for three consecutive days to induce differentiation, while the control group received PBS (**Fig. 4b**). Tumors were collected five days after bacterial injection, dissociated, and plated. Plated bacteria were resuspended in PBS and analyzed by flow cytometry to determine the proportions of each differentiated state (**Fig. 4c**).

Tumors from mice injected with the premixed consortium displayed a heterogeneous distribution of bacterial strains (**Figure 4d**). Furthermore, we observed that not all tumors were simultaneously colonized by all strains. Even in cases of triple colonization, the relative proportions of the populations varied significantly between tumors and largely deviated from the initial 1:1:1 ratio. These results align with our previous findings with a two-strain consortium, showing that injecting multiple strains results in heterogeneous colonization patterns across individuals, with tumors harboring different strain variant combinations and with different proportions.

Analysis of the differentiation groups revealed that, in the absence of induction, 97% of colonizing bacteria remained non-fluorescent, confirming the minimal *in vivo* leakage of our arabinose-inducible Bxb1 system and its long-term functionality (**Supplementary Fig. 18**). Following L-arabinose induction, all colonized tumors harbored a fully differentiated three-strain consortium, with relative abundances varying across individuals: 25–43% of S1, 42–56% of S2, and 4–36% of S3. Notably, the *in vivo* proportion of S2 bacteria closely mirrored the *in vitro* differentiation ratios (**Fig. 4d**, right panel). Interestingly, the S1 state was consistently overrepresented relative to the S3 strain, suggesting a fitness or initial-growth advantage for BFP-expressing cells in the tumor microenvironment. While the differentiation event establishes the initial ratios among consortia members, the final composition evolves according to the relative fitness of the daughter cells.

Mapping the relative proportions of S1, S2, and S3 from individual mice onto a ternary plot (**Fig. 4f**) revealed a clear contrast in population consistency between the two strategies. The co-administered mixtures (Group 1, dark diamonds) exhibited substantial inter-animal variability and deviated significantly from the initial 1:1:1 injection ratio (white dashed diamond). This scattering highlights the strong stochastic influence of the colonization bottleneck on independent strains. In contrast, the *in situ* differentiation system (Group 2, yellow circles) demonstrated a higher reliability. Population ratios were more tightly clustered and closer to the profile predicted from *in vitro* characterization (white dashed circle). These results indicate that our differentiation architecture effectively mitigates the founder effects that destabilize standard mixed-strain consortia, allowing the designed population distribution to be more reliably recapitulated within the tumor microenvironment.

Consequently, synthetic differentiation prevents single-strain dominance and consortium collapse, which are frequently observed in mixed approaches. By promoting a more balanced distribution of phenotypes within the microbial consortia, our system provides the functional diversity needed to achieve the predictable, reproducible outcomes required for effective bacterial therapies.

## DISCUSSION

Here, we engineered a multistate bacterial differentiation system using rationally designed competitive recombinase reactions. We show that the final ratio between daughter cells can be tuned by using recombination site variants, providing a robust mechanism for controlling the relative abundance of progeny phenotypes. Importantly, we show that the system is highly scalable and that higher-order states (up to 12 in this work) can be readily generated by sequentially adding additional recombination sites to the target DNA. In principle, this approach imposes no limit on the number of states, unlike earlier designs that rely on orthogonal dinucleotide attachment sites in series to create multiple states^35^. An essential characteristic of the system is that undifferentiated cells remain in a low-burden, off state during culture or transit, activating only upon reaching the target site, an essential feature for therapeutic bacteria^56^. This improves strain fitness and evolutionary stability, as well as the precision of payload delivery. A critical feature of our system is its reliance on a single recombinase and a single inducer, simplifying control over differentiation compared to other circuits that rely on multiple components^30,31^. This reduces cellular burden and avoids the complexities associated with balancing recombinases, directionality factors, and additional genes. Our design supports both excision and inversion, ensuring stable genomic encoding of genetic programs while enabling reversibility, if needed, through recombination directionality factors (RDFs). Finally, given that serine integrases do not require host-specific cofactors and work in a wide range of species^57,58^, this architecture could be adapted for mammalian systems to control differentiation ratios in cell-based therapies, such as defining the composition of CAR-T cell populations or controlling cell-type proportions in engineered tissues.

To demonstrate the system’s capacity to operate in complex physiological environments, we validated the functionality of our three-state differentiation system *in vivo*, where it outperformed traditional mixed-strain administration by reliably establishing consortia within tumor microenvironments and achieving reproducible, controllable ratios across individual mice. This reliability is essential for clinical translation, as colonization bottlenecks observed in transplanted tumor models are markedly exacerbated in more physiologically relevant settings. For example, in autochthonous mouse tumors, colonization efficiency can decrease by up to 10,000-fold relative to transplanted tumors, due to more restricted vascular access^59,60^. Similarly, early clinical trials reported limited bacterial tumor colonization in patients, likely due to tumor heterogeneity and more complex microenvironments^61–63^.

By decoupling phenotypic diversity from colonization efficiency, our *in situ* differentiation system provides a scalable platform for engineering functional diversification within the tumor microenvironment. For instance, one daughter strain could produce immunotherapeutic agents, while others would deliver cytotoxic payloads or modulate local metabolic balances, thereby enhancing therapeutic efficacy and minimizing systemic toxicity. Future work will determine whether this more robust establishment of phenotypic diversity translates into an improved therapeutic activity compared to the standard “mixture of strains” strategy. Other applications include gut microbial intervention, in which bacteria equipped with differentiation systems responsive to specific localization, pathological, or external signals could generate microbial communities tailored to distinct microenvironments. Such an approach would enable targeted restoration of dysbiotic microbiomes, treatment of metabolic disorders, and immune modulation, with potential applications in inflammatory bowel disease and colorectal cancer prevention. Beyond therapeutics, our differentiation systems hold promise for bioproduction, where a division of labor among strains can optimize yields of biofuels, pharmaceuticals, or specialty chemicals^64^. For bioremediation, progenitor strains could differentiate *in situ* into specialized daughter cells, each activating distinct dormant detoxification pathways in response to environmental signals, enabling the efficient degradation of complex pollutants and the remediation of contaminated sites^65,66^.

We found that the DNA distance between the first two sites strongly influences site preference during competitive excision. We identified a biophysical threshold below ∼200 bp where recombination was highly biased toward distal sites. We attribute this effect to the persistence length of double-stranded DNA (∼150 bp)^45^, which imposes a high energetic penalty on the formation of the compact synaptic loops required for proximal recombination. While these physical limitations are less apparent in non-competitive contexts^67^, the recombinase’s ability to select the energetically favorable path highlights these differences in our system. We also observed that once this minimal length threshold was crossed, the preference reversed, favoring the shorter distance. This indicates a transition from a stiffness-limited regime to a diffusive search regime, in which the higher effective concentration of proximal sites dictates the recombination frequency (see **Supplementary Note 1** for extended discussion). Interestingly, maximizing the proportion of cells in State 1 (S1) for the three-state system requires combining fast-kinetic sites with increased inter-site spacing (**Fig. 2d**).

A common challenge in engineering microbial consortia is maintaining population stability, as metabolic burden often drives the competitive exclusion of slower-growing strains. Indeed, we observed that in well-mixed *in vitro* batch cultures, slight differences in fitness between differentiated subpopulations led to gradual drift in community composition over time. However, this instability was attenuated in the *in vivo* tumor environment, where consortia compositions remained relatively robust across individuals two days post-differentiation. We hypothesize that the spatial structure of the tumor microenvironment imposes physical constraints that buffer against competitive exclusion, preventing the rapid takeover of fast-growing clones observed in liquid culture. Furthermore, the tunability of our differentiation architecture offers a promising design strategy to counteract metabolic drift. Since therapeutic payloads inevitably impose varying physiological burdens, our system allows the initial differentiation ratio to be biased in favor of the metabolically burdened strain. By starting with a larger pool of the slower-growing subpopulation, the onset of competitive exclusion could be delayed, thereby extending the operational lifespan of the engineered consortium before equilibrium shifts render it nonfunctional. To further enhance stability, future iterations could integrate metabolic interdependencies (such as obligate cross-feeding) between subpopulations, reducing the potential for dominance by any single phenotype. Additionally, fine-tuned modulation of gene expression levels could be employed to balance fitness costs across daughter cell types, ensuring a more uniform growth dynamics.

In conclusion, our differentiation circuits provide a simple, modular, and scalable strategy for controlling differentiated cell ratios using a single input. By enabling precise modulation of phenotypic diversity in microbial consortia, our work provides a robust foundation for advancing applications in synthetic ecology, bioproduction, and programmable therapeutics.

## MATERIALS AND METHODS

### Bacterial strains and culture medium

All bacterial cloning experiments were performed using *E. coli NEB 10 Beta* electro-competent cells prepared in-house. Differentiation experiments were performed in *E. coli NEB 10 Beta* and *E. coli Nissle 1917. E. coli* cultures were grown in lysogeny broth (LB) medium with appropriate antibiotics, at 30°C or 37°C with shaking at 250 rpm. To maintain plasmids, LB medium was supplemented with the following antibiotics: chloramphenicol at 25 µg/mL, carbenicillin at 100 µg/mL, kanamycin at 50 µg/mL, and spectinomycin at 100 µg/mL. L-arabinose (Sigma-Aldrich, Cat. No. A3256) was used to a final concentration of 0.1% w∕vol. For the co-transformation of two plasmids, the two corresponding antibiotics were used at the previously defined concentration.

### Plasmid constructions

All plasmids used in this study are described in **Supplementary Table 1**. Cloning was either performed by GenScript (Rijswijk, Netherlands) or in-house using gene fragments synthesized by IDT (Leuven, Belgium) or Twist Biosciences (South San Francisco, USA). We used the provider’s algorithm to codon-optimize genes for *E. coli.* DNA assembly was performed using the Gibson assembly method (Gibson et al. 2009) with vector backbones pSB4K5 (Shetty et al. 2008), pBAC (Wild et al. 2002), or pDonor (Vo et al. 2021). The constitutive Bxb1 expression plasmid is from Guiziou *et al.*^39^ (Addgene #117030), and the plasmid with p15Aori containing the arabinose-inducible Bxb1cassette is from Bonnet *et al.*^8^ (Addgene #38209).

### DNA transformation into *E. coli NEB 10 Beta* and inoculation

SOC medium was pre-warmed at 37°C. Electro-competent *E. coli NEB 10 Beta* were thawed on ice. 100 ng of each plasmid was gently mixed with 50 µL of electro-competent *E. coli* NEB *10 Beta*. Co-electroporation was performed of a constitutive Bxb1 plasmid and a BAC or pSB4K5 differentiation circuit with Bio-Rad MicroPulser with a 0.1 cm cuvette (bacteria 1.8 kV), with 100 ng/µL of each plasmid. Then, 900 µL of SOC was immediately added to the cuvette, and the cells were resuspended and transferred to a 1.5 mL culture tube. Recover for one hour at 37°C, shaking at 190 rpm. 5 µL of recovery was inoculated into 495 µL of LB medium containing the appropriate antibiotics (dilution 1:100) in a 96-deep well, with technical replicates. The deep well was incubated in a Kuhner with shaking at 250 rpm overnight. For the differentiation experiments using sfGFP and mKate2, cultures were incubated overnight at 30°C. In contrast, experiments involving BFP were carried out overnight at 37°C, as BFP requires a longer maturation time^50^. At 30°C, a substantial fraction of cells remained undifferentiated after an overnight incubation in this case.

### Small-batch electro-competent cells preparation and transformation

For the preparation of small-batch electro-competent cells, a single colony was inoculated into LB medium and grown overnight at 37°C and 190 rpm. The following morning, 300 µL of the overnight culture was transferred into 10 mL of fresh LB in a 50 mL Falcon tube and incubated at 37°C, 190 rpm until the culture reached an OD_600_ of ∼0.5-0.6. 1 mL of culture was transferred into a 1.5 mL tube and centrifuged at 11,000xg for 1 min at 4°C. The cell pellet was washed three times with 1 mL of ice-cold 10% sterile glycerol, with centrifugation at 11,000xg for 1 min at 4°C. After each wash, the pellet was resuspended to ensure efficient removal of residual medium. Following the final wash, the pellet was resuspended in 50 µL of 10% glycerol and immediately used for electroporation with the plasmid of interest.

### Colony-forming units (CFU)

Bacterial cultures were placed in the first row of a 96-well plate in 3 technical replicates. The first row is then diluted 1/10 in sterile PBS, and this dilution is carried out serially through row 8. A 1 µL drop from each well is then transferred using a 6x8 replica plater (Sigma-Aldrich) onto an LB agar petri dish and allowed to dry. Following overnight incubation of the petri dish at 37°C, CFUs were counted in the first countable dilution drop from each replicate, and the bacterial concentration was calculated from the dilution.

### Integrate transposition experiments

Differentiation circuits were integrated into the *E. coli* chromosome using a CRISPR RNA–guided transposon system (Tn6677 from *Vibrio cholerae*)^68^. Electrocompetent cells were co-transformed with a donor plasmid carrying the differentiation circuits (pDonor) and an effector plasmid encoding the transposition machinery and the crRNA targeting the desired chromosomal locus (pEffector). Following electroporation, cells were allowed to recover for one hour at 30°C. Transformants were then selected on LB agar supplemented with ampicillin and spectinomycin at 30°C for 24h. The next day, colonies were collected by adding 5ml of LB directly onto the plate, scraping all colonies with a sterile glass pipette, and homogenizing the suspension. A 1:10,000 dilution was prepared (1µl into 999µl LB, followed by 10µl into 990µl LB), and 100µl was replated on fresh ampicillin- and spectinomycin-containing LB agar, at 30°C for 24h.

In our experiments, two different chromosomal loci were targeted: the *YFC* locus in the *E. coli 10 Beta* strain and the *lacZ* locus in the *E. coli Nissle* strain. For the *YFC* locus, individual clones were screened by colony PCR to confirm correct chromosomal integration. Colonies were resuspended in 30µl sterile water and used directly as templates. PCR was performed using Phusion DNA polymerase with primers described in Supplementary Table n°1. Products were resolved on 0.8% w∕vol agarose gels with a 100bp DNA ladder.

For integration in the *lacZ* locus, after the first plating, colonies were then pooled by resuspension in LB, diluted 1:1,000, and replated on LB agar containing ampicillin, spectinomycin, isopropyl β-D-1-thiogalactopyranoside (IPTG; 1M; Euromedex), and 5-bromo-4-chloro-3-indolyl-β-D-galactopyranoside (X-gal; 0.1 mg mL⁻¹; Sigma-Aldrich). After overnight incubation at 30°C, individual colonies were first screened by blue–white color selection on X-Gal/IPTG plates. White colonies, corresponding to disruption of the *lacZ* gene and therefore indicative of successful chromosomal integration at the target locus, were selected for further analysis. Chromosomal integration was subsequently confirmed by colony PCR as described below.

### Plasmid Curing

To cure the remaining plasmids after chromosomal integration, the pFREE system was used^50,69^. Briefly, strains harboring the plasmids were grown overnight in 3ml LB containing the appropriate antibiotics. The following day, electrocompetent cells were prepared by inoculating 10ml LB with 300µl of the preculture and transformed with 100ng of pFREE using 50µl of competent cells. After recovery for two hours at 30°C in 900µl SOC with shaking at 190 rpm, 10µl of the culture was transferred into 10ml LB supplemented with 0.2% L-rhamnose w∕vol (Sigma-Aldrich, Cat. No. W373011), 200ngml⁻¹ anhydrotetracycline (aTc), and 50µgml⁻¹ kanamycin to induce gRNA-guided Cas9 cleavage of the target plasmids. Cultures were incubated for 24h at 30°C with agitation. Cells were then plated on non-selective LB agar, diluted by bead transfer, and incubated for 24h at 30°C. Plasmid curing was assessed by colony PCR targeting plasmid replicons and by phenotypic screening for antibiotic sensitivity.

### Genomic DNA extraction and whole-genome sequencing

Single bacterial colonies were inoculated into 10 ml of LB medium supplemented with the appropriate antibiotics and grown overnight at 37°C with shaking at 190 rpm. Genomic DNA was extracted from the saturated culture using a commercial genomic DNA purification kit (Quick-DNA Miniprep Plus Kit, Zymo Research, Cat. No. D4069) according to the manufacturer’s instructions, including RNase treatment to remove residual RNA. DNA concentration was measured, and purity was assessed by spectrophotometry (NanoDrop, Thermo Fisher Scientific).

### Flow cytometry

For flow cytometry, stationary cultures were diluted 1:100 into 200µl of 1X PBS in flat-bottom 96-well plates. Fluorescence measurements were acquired on an Attune NxT Flow Cytometer (Thermo Fisher Scientific). The following settings were used: forward scatter (FSC) 350V with a 488nm laser, side scatter (SSC) 380V with a 488nm laser, green fluorescence (BL1 channel, sfGFP) 520V with 488nm excitation, red fluorescence (YL2 channel, mKate2) 550V with 561nm excitation, and blue fluorescence (VL1 channel, BFP) 410V with 405nm excitation. For each sample, data from 30,000 events were collected. For each sample, data from 30,000 events were collected. All experiments included a negative control consisting of cells lacking a plasmid and grown under identical conditions, and a positive control expressing either sfGFP, mKate2, or BFP, which was used to define fluorescence gates. Data were analyzed using FlowJo (Treestar, Inc.). Each experiment was performed in biological triplicate, consisting of three independent transformations conducted on different days. For each experiment, three technical replicates were included in 96-well plates.

### Growth curve measurements

Bacterial growth was monitored by measuring absorbance at 600nm using a Biotek Cytation 3 or Biotek Synergy H1M microplate reader. Glycerol stocks were streaked on LB agar plates containing the appropriate antibiotics and incubated overnight at 37°C. Single colonies were used to inoculate 96-deep-well plates containing 500µl of LB medium supplemented with the appropriate antibiotics, and the plates were incubated overnight at 37°C with shaking at 250rpm in biological triplicate. The following day, experimental cultures were prepared by inoculating 198µl of fresh LB medium (with or without antibiotics and with or without inducers, including 0.1% (w/v) L-arabinose for Bxb1 induction) with 2µl of a preculture previously diluted 1:1,000 in fresh medium, in flat-bottom 96-well plates. Plates were incubated in the plate reader preheated to 37°C with continuous orbital shaking between measurements. Absorbance at 600 nm was recorded every 10min overnight, and wells containing LB medium only were used for background subtraction. When fluorescence was measured simultaneously, excitation/emission settings were as follows: BFP (375nm/440nm), sfGFP (485nm/528nm), and mKate2 (585nm/630nm), with a gain of 100.

### Co-culture passaging

To evaluate the long-term stability of bacterial consortia, *E. coli* strains expressing either sfGFP or mKate2 were mixed at a 50:50 ratio and cultured in 500 µL of LB medium supplemented with appropriate antibiotics at 37 °C with shaking at 250 rpm overnight. In the morning, saturated cultures were analyzed by flow cytometry. For each subsequent round of growth, cultures were diluted 1:250 into 500 µL of fresh LB medium with appropriate antibiotics and incubated again overnight under the same conditions. This cycle of growth, dilution, and flow cytometry analysis was repeated for 5 days.

To assess the long-term stability of genetic circuits in the absence of inducers, bacterial cultures were diluted 1:250 every 12 days. At each passage, cells were analyzed by flow cytometry to detect spontaneous differentiation events resulting from potential recombinase leakiness.

### Confocal fluorescence microscopy on agar pad

Bacterial cultures were grown overnight in LB medium with the appropriate antibiotics at 37 °C with shaking at 190 rpm. For imaging, 2% (w/v) low–melting–point agarose was poured as a thin layer onto glass slides or into custom casting molds to form flat agarose pads. Once solidified, pads were cut to size and placed on glass microscope slides. 1 µL of the bacterial culture was spotted onto the agarose pad and allowed to absorb for 1–2 min. Pads were then gently covered with a clean coverslip. Images were acquired using an EVOS M7000 digital inverted microscope (Thermo Fisher Scientific) equipped with a 60× oil-immersion objective (Olympus). On the EVOS M7000, BFP was imaged using the DAPI LED cube (excitation/emission: 357/447 nm), sfGFP using the GFP LED cube (482/524 nm), and mKate2 using the RFP/Texas Red LED cube (542/593 nm or 585/628 nm). All images were processed using ImageJ software (version 1.54c, NIH) under identical settings for each experiment.

### Long-read sequencing and analysis

Sequencing libraries were prepared from purified DNA using the Rapid Sequencing Kit (SQK-RBK114.96 Oxford Nanopore Technologies) according to the manufacturer’s instructions. DNA samples were subjected to tagmentation with a transposase, enabling simultaneous fragmentation and attachment of sequencing adapters. The rapid adapter mix was then added, followed by a 5-min incubation at room temperature. Libraries were immediately loaded on a MinIon R10.4.1 fongle flow cell (FLO-FLG114). Long-read sequencing was performed on a MinION Mk1B (Oxford Nanopore Technologies). Runs were conducted for 12 to 24 h, depending on pore activity and the available starting material. Basecalling was performed using Guppy (v6.5.7) in Super Accuracy (SUP) mode.

Sequencing reads from each biological replicate were mapped to the reference genome using minimap2 (v2.24) with default parameters^70^. The resulting SAM files were converted to BAM format, sorted, and indexed using pysam (v0.19.1). Only aligned reads were retained for downstream analysis. Recombination breakpoints were identified based on the presence and position of *attB* recombination sites in the reference genome. The *attB* sequence (5’-GCCCGGATGATCCTGACGACGGAGACCGCCGTCGTCGACAAGCCGGCCGA-3’, 51 bp) was automatically detected using Biopython’s SeqIO module^71^. Breakpoint positions were calculated as *attB* start position + 21 nucleotides, corresponding to the recombinase cut site between the two guanine nucleotides within the *attB* sequence. Reads were classified into distinct categories based on their start positions relative to detected breakpoints. Reads mapping upstream of the first breakpoint were classified as non-recombinant (category Break0), while reads mapping between successive breakpoints were classified as recombinant (categories Break1 to BreakN, where N is the number of intervals between breakpoints). The percentage of reads in each category was calculated relative to the total number of aligned reads. Mean percentages and standard deviations were computed across replicates using NumPy (v1.21.0), with standard deviations calculated using Bessel’s correction (ddof=1). Read distributions along the reference genome were visualized as histograms with 1000 bins using Matplotlib (v3.5.0). Breakpoint positions were marked with vertical lines, and intervals between breakpoints were color-coded to distinguish recombinant categories. Bar plots showing the percentage of reads per category were generated for individual replicates and for mean values across all replicates, with error bars representing standard deviations. To validate breakpoint positions, read start position peaks were independently detected using scipy’s find_peaks function (v1.7.0) with a prominence threshold of 10. Detected peaks were compared to calculated breakpoint positions for quality control.

The script that automates the entire workflow and the raw sequencing files are available on the GitHub repository: https://github.com/asfistonlavie/RecombCompet/.

### Mice tumor-bearing model

Mice experiments were performed in accordance with protocols approved by the Languedoc-Roussillon Animal Ethics Committee (Comité d’Éthique en Expérimentation Animale Languedoc-Roussillon), an institution accredited by the French Ministry of Higher Education, Research and Innovation. All experiments were conducted using 6–8-week-old female C57BL/6J mice (Charles River Laboratories), which were housed in a pathogen-free barrier facility (room temperature: 22 °C; relative humidity: 55%; 12-h light/dark cycle). Animal housing and procedures, including euthanasia, were carried out in accordance with the 3R rules. Ectopic colorectal tumors were grafted onto the flank of the mice by subcutaneous injection of 200 µL containing 5 x 10^5^ murine colon adenocarcinoma cells (MC38). The MC38 certified cell line was kindly provided by the Integrated Research Center on Cancer (SIRIC) and was grown in Dulbecco’s modified MEM supplemented with 10% fetal bovine serum.

Tumors were measured using an electronic caliper, and volumes were calculated from their length and width (V = L * W2/2). Tumors were grown to an approximate volume of 200 mm^3^ before randomizing the mice into groups with the same mean tumor volume and tumor growth rate right before the bacterial intravenous injection. Animals were sacrificed by cervical dislocation five days after bacterial injection.

### Bacterial administration

Bacteria were counted by flow cytometry and adjusted to the desired concentration. Briefly, an overnight-grown EcN culture was washed 3 times with ice-cold, nutrient-free PBS. Then, a counting dilution was prepared to obtain 1 million bacteria in 100 µL for counting with the flow cytometer. Counting was performed on an Attune NxT flow cytometer equipped with a KickMax autosampler and Attune NxT Version 2.7 Software (Thermo Fisher). The actual concentration of the washed solution was then calculated back from the dilution absolute count. After this first count, the solution was diluted or concentrated to match the targeted concentration. Bacteria were injected intravenously via the tail vein of the mice at a total volume of 100 µL. Following the injection, the remaining bacteria were plated to assess the injected concentration further by CFU counting.

### In vivo imaging

*E. coli Nissle 1917* expressing bioluminescence (a kind gift of Dr. L.A. Fernandez, CNB-CSIC) was detected *in vivo* using an IVIS Lumina III spectrum imaging system and Living Image® software (PerkinElmer). The total photons per second detected were used to quantify the bioluminescence emitted by *EcN* in identical-sized gates for each mouse.

### Biodistribution

At the study endpoint, mice were euthanized by cervical dislocation. The tumor and selected organs were collected, weighted and homogenized using the gentleMACS tissue dissociator (Miltenyi Biotec; C-tubes) in 8 ml PBS containing 50 µL collagenase B and 5 µL DNase I. Homogenates were filtered through 70 µm cell strainers (Corning), serial diluted in PBS and plated on LB agar plates as described in the CFU count section to quantify the bacterial colonization. CFU were counted the next day, and colony bioluminescence was assessed using the Amersham imager 600 (GE Healthcare Life Sciences) to confirm that the retrieved bacteria corresponded to the injected *EcN-lux*.

### Statistics and reproducibility

All data were processed using GraphPad Prism, Microsoft Excel, or in-house scripts. Error bars in the figures represent standard deviation, as specified in the figure legends.

## ACKNOWLEDGEMENTS

We thank members of our groups and institutes for fruitful discussions and feedback. We thank Mikhail Shapiro (Caltech) and Luis Angel Fernandez (CNB-CSIC) for kindly providing the *ARG* and *luxCDABE* operons, respectively. We thank Isabelle Teulon, Adeline Toro, and the members of the IRCM’s animal facility unit (BioCampus RAM-PEFO, IRCM, Montpellier, France) for animal care and *in vivo* studies. We thank Sarah Guiziou (Earlam Institute), David Maresca (TU Delft), Celine Deraison (IRSD), Celine Gongora (IRCM), and William Bourguet (CBS) for constructive feedback on this work. We thank Lukas Brichet of SMARTLife Biosciences for assistance with long-read sequencing of the samples. We are also grateful to Laurine Dal Toe for her help with ImageJ analyses. JB thanks INSERM and the Bettencourt-Schueller Foundation for their continuous support. This work was supported by an ANR grant “SonoGT” (ANR-21-CE19-0050), a grant from PEPR “BBEST”, and by an INSERM grant for the development of a Technological Research Accelerator in Synthetic Biology (ART-synbio). CS is a recipient of a 4th-year PhD fellowship from the ARC Foundation. QB, EF, and JuB are recipients of a PhD fellowship from the French Ministry of Research. The CBS acknowledges support from the French Infrastructure for Integrated Structural Biology (FRISBI; ANR-10-INSB-05-01).

## Authors contributions

JB and CS conceived the study. CS conducted gene construction, flow cytometry, and microscopy experiments. CS, JC, and AAK performed chromosomal integrations. MG, CS, AAK, QB, EF, and JuB conducted mouse experiments. CS, JC, AAK, and JB analysed data. CM and ASFL performed long-read analysis. XD performed data processing and visualization. JB, MCG, and DC supervised the study. JB and CS wrote the manuscript with input from all authors. All authors read and approved the final version of the manuscript.

## Competing interests

The authors declare a competing interest in the form of patent applications related to the synthetic differentiation designs presented in this study.

